# Macrophage-Derived 25-Hydroxycholesterol Drives Statin-Associated Acute Kidney Injury

**DOI:** 10.64898/2026.10.16.759612

**Authors:** Hye Jin Wang, Eun Jee Kim, Yuri Cho, Jung Ho Lee, Jongjin Yoon, Hye-Youn Kim, Youngrak Jung, Haebeen Lee, Hoon Young Choi, June-Yong Lee, Heon Yung Gee, Chul Hoon Kim, Beom Seok Kim

## Abstract

Although acute kidney injury (AKI) has been implicated as a potential adverse effect of statin therapy, experimental evidence supporting this association remains limited. Here, we investigated whether elevated levels of 25-hydroxycholesterol (25-HC), an oxysterol produced by activated macrophages and monocytes, increase susceptibility to statin-induced AKI. We found that atorvastatin and fluvastatin induced cell death in proximal tubular cells when combined with 25-HC. Consistently, administration of low-dose lipopolysaccharide triggered AKI in statin-treated mice. Mechanistically, impaired protein prenylation disrupted the autophagosome–lysosome pathway in proximal tubular cells. AKI was attenuated by mevalonate supplementation, which restored protein prenylation, and by macrophage/monocyte depletion, which reduced the cellular sources of 25-HC. Furthermore, combined exposure to fluvastatin and 25-HC increased kidney injury molecule-1 (KIM-1) expression in human induced pluripotent stem cell-derived kidney organoids. Together, these findings suggest that elevated macrophage-derived 25-HC under conditions of infection, inflammation, or major surgery may increase susceptibility to statin-induced AKI. Our study identifies a previously unrecognized interaction between statins and 25-HC and suggests potential strategies to mitigate kidney injury while preserving the therapeutic benefits of statins.

## INTRODUCTION

Statins are a class of lipid-lowering drugs that reduce circulating levels of low-density lipoprotein (LDL) cholesterol by inhibiting 3-hydroxy-3-methylglutaryl–coenzyme A (HMG-CoA) reductase, the rate-limiting enzyme in hepatic cholesterol biosynthesis. This inhibition decreases intracellular cholesterol production in the liver, leading to upregulation of LDL receptors and enhanced clearance of LDL cholesterol from the bloodstream (1). Because they effectively decrease the risk of atherosclerotic cardiovascular disease (ASCVD) and stroke, statins are among the most widely prescribed medications worldwide (2). Statin therapy, like all pharmacologic treatments, is associated with adverse effects. For instance, statins can increase blood glucose levels, which may contribute to the development of type 2 diabetes (3, 4). In rare cases, particularly at high doses, statins can cause muscle cell breakdown (rhabdomyolysis), leading to the release of myoglobin into the bloodstream and resulting in severe muscle pain and possible kidney damage (5). Nevertheless, for most patients, the cardiovascular benefits of statin therapy substantially outweigh these relatively limited risks (2, 4).

More recently, statin use has also been linked to an increased risk of acute kidney injury (AKI). AKI is characterized by an abrupt decline in glomerular filtration rate (GFR) sufficient to impair the elimination of nitrogenous waste products and other uremic toxins. It is a common and clinically significant complication among hospitalized patients across a wide range of conditions (6). Although AKI can be reversible, it is associated with substantial morbidity and mortality, and no definitive treatment currently exists (7). Notably, two large observational studies have suggested that statin therapy may increase the risk of AKI (8, 9). Furthermore, a more recent randomized clinical trial (RCT) reported that high-dose perioperative atorvastatin was associated with higher rates of AKI in patients with underlying chronic kidney disease (CKD) who were statin-naïve prior to study enrollment (10).

Similarly, another RCT found that perioperative rosuvastatin use in patients undergoing cardiac surgery increased the incidence of postoperative AKI (11). In contrast, several other studies have suggested that statins may confer protective effects against postoperative AKI (12–14). These conflicting findings indicate that statins may either protect against or contribute to kidney injury depending on clinical context and patient characteristics. Thus, whether statins truly increase the risk of AKI—and, if so, which factors determine individual susceptibility to statin-induced AKI—remains an unresolved question, warranting well-designed preclinical model studies that enable rigorous control of experimental variables to address this issue. Given that even mild AKI is associated with a significantly increased risk of serious adverse outcomes (15), identifying the molecular determinants of susceptibility to statin-associated AKI is of considerable clinical importance.

In this study, we hypothesized that statins may induce AKI by directly affecting renal cells under specific pathological conditions, such as infection or inflammation. This hypothesis is grounded in the observation that statins induce AKI in only a small subset of cases and are more often implicated in settings where additional stressors—particularly CKD or major surgery—are present (10, 11), suggesting these specific conditions may underlie the association. We first performed an *in vitro* screen to identify immune and inflammatory mediators that interact with statins and influence tubular cell viability. This approach identified 25-hydroxycholesterol (25-HC) as a candidate mediator. We then validated this finding *in vivo* by demonstrating that lipopolysaccharide (LPS) stimulation, a condition known to elevate 25-HC levels, induced AKI in statin-treated mice. Notably, both macrophage/monocyte depletion and genetic ablation of the 25-HC–synthesizing enzyme in these cells prevented the onset of AKI, underscoring that these immune cells constitute a critical source of 25-HC and thereby act as key mediators of this statin-associated renal injury. Furthermore, we reproduced the adverse interaction between statins and 25-HC in human induced pluripotent stem cell (iPSC)–derived kidney organoids. Finally, we propose that renal cell damage induced by combined statin and 25-HC exposure is mediated by impaired protein prenylation and consequent disruption of autophagosome–lysosome function.

## MATERIALS and METHODS

### Animals

All mice were maintained under specific pathogen-free conditions at room temperature and 40%–60% humidity, with free access to food and water. The mice were allowed to acclimatize for 7 days before the start of the experiments

### Statin-induced AKI mouse model and *in vivo* study design

C57BL/6J mice, GFP-LC3 transgenic mice, and homozygous *Ch25h* knockout (*Ch25h*^-/-^) mice with littermate wild-type (*Ch25h*^+/+^) controls were placed on a 60% high-fat diet (HFD) beginning on experimental day 0. Atorvastatin (1 mg/kg, intraperitoneally [IP]) was administered daily during days 7 to 10. To induce inflammation, mice received LPS (2 mg/kg, IP) on day 9. Blood samples were collected 30 h after LPS administration, and mice were sacrificed 30 or 48 h post-LPS treatment for histological analyses. For immunostaining of NGAL and cleaved caspase-3, a separate cohort of mice was fed a chow diet containing fluvastatin (25 mg/kg/day) for 2 weeks. On day 10, the mice received LPS (2 mg/kg, IP). Kidneys were harvested 3 days later and fixed for cryosectioning.

For the rescue experiments, mice were fed a 60% HFD from experimental day 0. They were given both atorvastatin (1 mg/kg, IP) and mevalonolactone (10 mg/kg, IP) daily from day 7 to day 10. The mice were injected with LPS on day 9. Blood and kidney samples were collected 30 h after LPS administration for the assessment of blood BUN and creatinine levels, as well as for histological analysis.

### *In vitro* modeling of statin-induced AKI using human kidney organoids

To induce and evaluate AKI-related pathology *in vitro*, fully differentiated human kidney organoids were cultured for 72 h in KR medium supplemented with 25-HC (0.1 μg/mL), either alone or in combination with fluvastatin (5 μM) or atorvastatin (5 μM). Following treatment, the organoids were harvested and processed for downstream pathological analyses, including immunofluorescence staining for proximal tubular injury markers and transmission electron microscopy (TEM).

## RESULTS

### Statins induce 25-HC-dependent HK-2 cell toxicity through impaired protein prenylation *in vitro*

Statins are generally safe but have been associated with AKI in specific clinical settings, including CKD and major surgery (10, 11), where infection or inflammation is common (16–18). We therefore tested whether inflammatory or immune mediators cooperate with statins to induce toxicity in HK-2 proximal tubular cells. Among the mediators examined, only 25-HC enhanced atorvastatin-induced LDH release, whereas neither 25-HC nor atorvastatin alone caused significant cell death (Fig. 1A–C). 25-HC is a macrophage-derived oxysterol involved in lipid metabolism and inflammatory responses (19, 20). This toxic interaction was reproduced with fluvastatin and was also observed with other major oxysterols, including 24-HC and 27-HC (Fig. 1B–D). In the plasma, oxysterols are transported mainly by lipoproteins (21). Oxysterol-enriched LDL similarly sensitized HK-2 cells to statin-induced death, and the combination of statins and 25-HC showed selective toxicity in primary human proximal tubular cells, but not in mesangial cells or mouse hepatocytes (Supplementary Figs. 1 and 2).

**Fig. 1.**
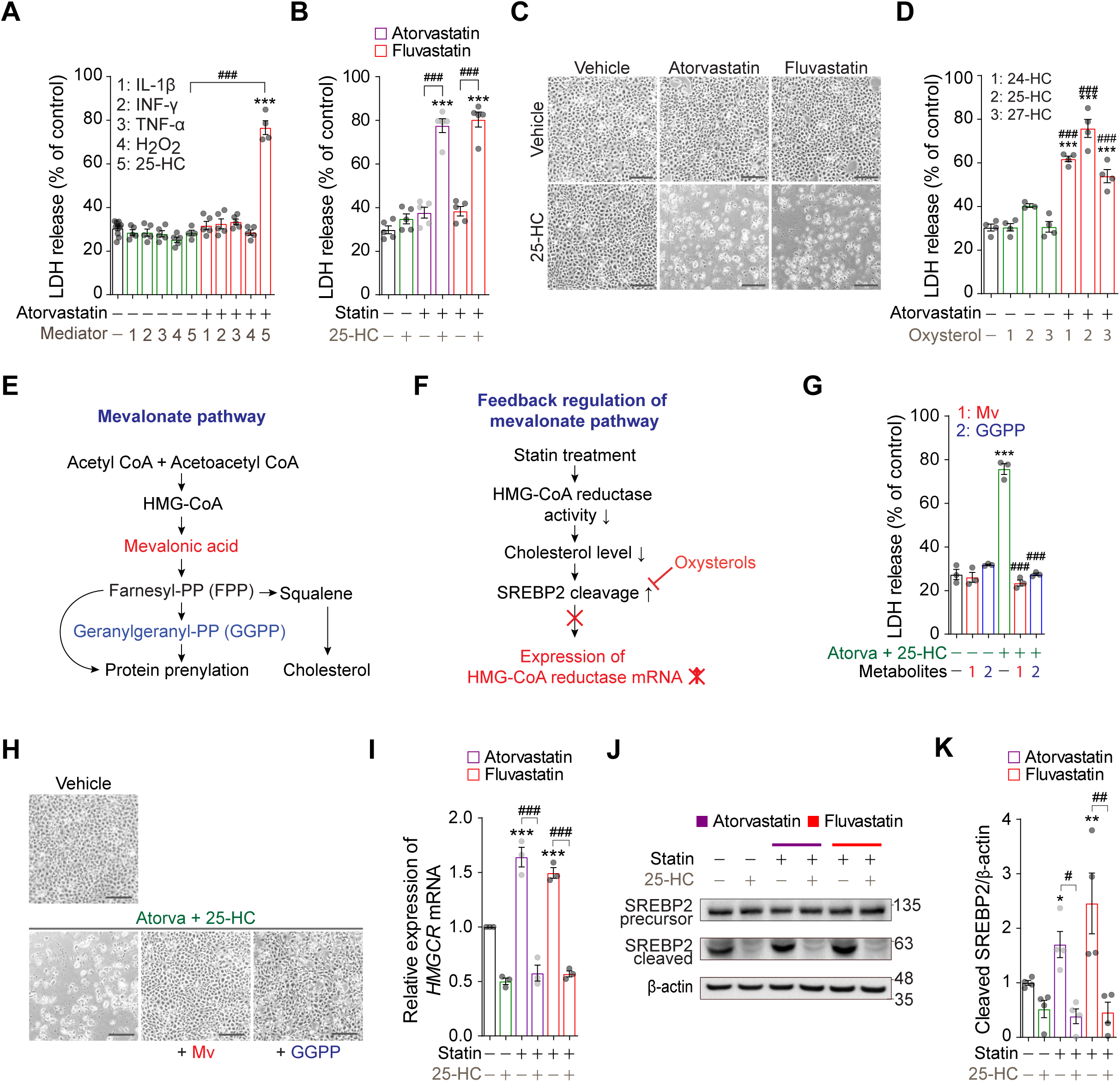
Statins induce 25-HC-dependent HK-2 cell toxicity through impaired protein prenylation *in vitro*. (**A**) HK-2 cells were treated for 72 h with atorvastatin (5 μM) and an inflammatory mediator (IL-1β, IFN-γ, or TNF-α, 0.01 ng/ml each; H₂O₂, 1 μM; or 25-HC, 0.1 μg/ml). Cell death was assessed by measuring LDH release into the culture medium^. ***^*P* < 0.001 vs. untreated control; ^###^*P* < 0.001 vs. 25-HC alone. (**B**) HK-2 cells were treated for 72 h with atorvastatin (5 μM) or fluvastatin (1 μM), with or without 25-HC (0.1 μg/ml). Cell death was assessed by LDH assay. \*\*\**P* < 0.001 vs. untreated control; ^###^*P* < 0.001 vs. statin alone. (**C**) Representative bright-field images showing HK-2 cell death after 72 h of combined statin and 25-HC treatment. Scale bars: 250 μm. (**D**) HK-2 cells were treated for 72 h with atorvastatin (5 μM) and an oxysterol (24-HC, 25-HC, or 27-HC; 0.1 μg/ml each). Cell death was assessed by LDH assay. \*\*\**P* < 0.001 vs. untreated control; ^###^*P* < 0.001 vs. each oxysterol alone. (**E**) Schematic of SREBP2-mediated feedback activation leading to upregulation of HMG-CoA reductase expression. (**F**) Schematic illustration of the mevalonate pathway involved in isoprenoid and cholesterol synthesis. (**G**) HK-2 cells were treated for 72 h with atorvastatin (Atorva, 5 μM) and 25-HC (0.1 μg/ml) in the presence or absence of mevalonate pathway metabolites, including mevalonolactone (a lactone form of mevalonic acid; 200 μM) or geranylgeranyl pyrophosphate (GGPP, 10 μM). Cell death was quantified using the LDH assay. \*\*\**P* < 0.001 vs. untreated control; ^###^*P* < 0.001 vs. atorvastatin plus 25-HC treatment. (**H**) Representative bright-field images demonstrating that supplementation with mevalonolactone (200 μM) or GGPP (10 μM) rescues HK-2 cells from combined treatment-induced cell death. Scale bars: 250 μm. (**I**) HK-2 cells were treated for 24 h with statins (atorvastatin, 5 μM; or fluvastatin, 1 μM) in the presence or absence of 25-HC (0.1 μg/ml), followed by real-time PCR analysis. \*\*\**P* < 0.001 vs. untreated control; ^###^*P* < 0.001 vs. statin treatment alone. (**J**) HK-2 cells treated with statins and 25-HC for 12 h were subjected to western blot analysis using anti-SREBP2 antibodies. (**K**) Quantification of cleaved SREBP2 band intensities normalized to the corresponding actin band intensities. \**P* < 0.05 and \*\**P* < 0.01 vs. untreated control; ^#^*P* < 0.05 and ^##^*P* < 0.01 vs. statin treatment alone. Data in (**A**, **B**, **D**, **G**, **I**, and **K**) are presented as mean ± SEM, with individual data points shown (n ≥ 3 per group).

**Fig. 2.**
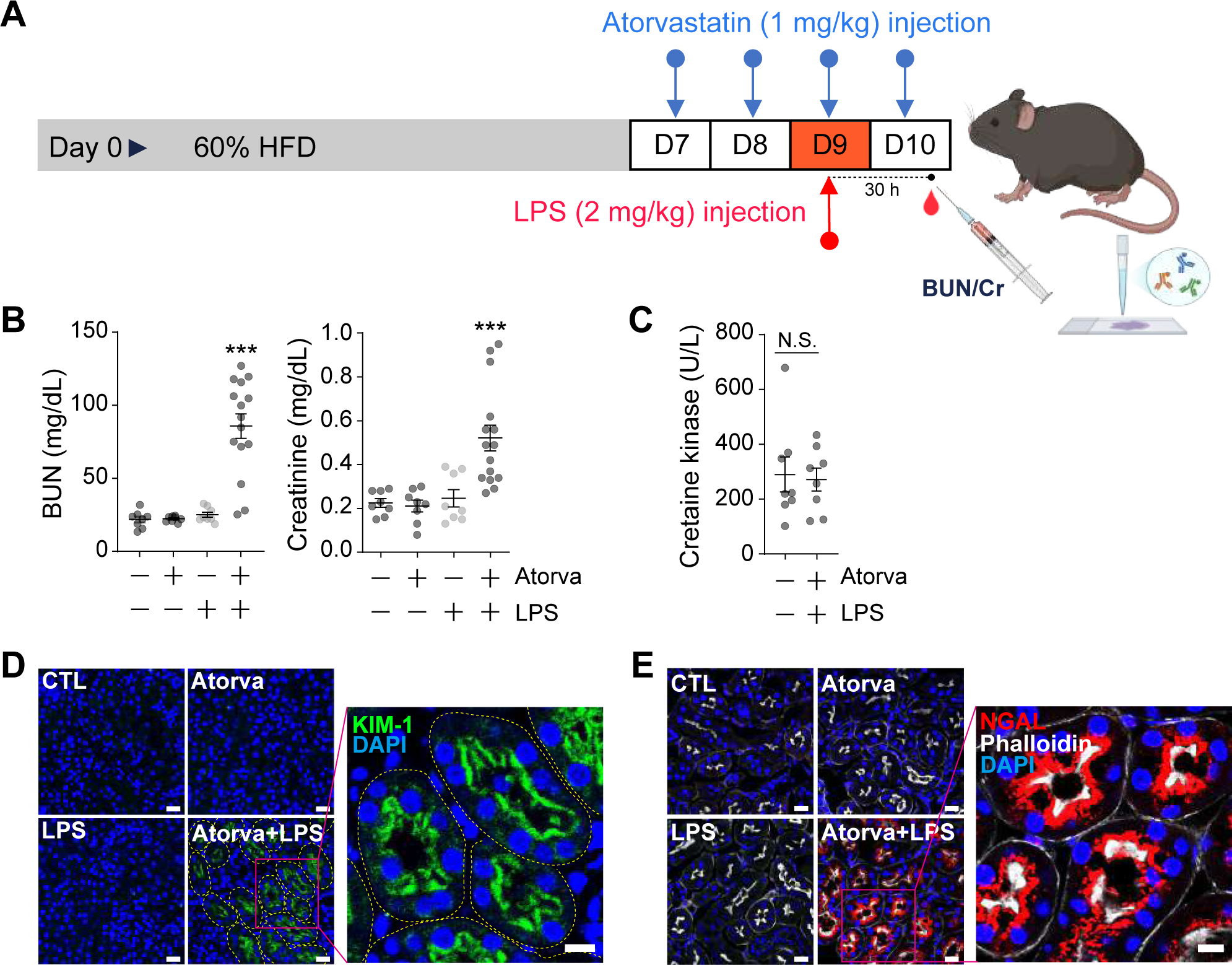
Atorvastatin-induced AKI in an LPS-induced inflammatory mouse model. (**A**) Schematic of AKI induction in mice. Mice were fed a 60% HFD daily from day 0 until the end of the experiment, with daily atorvastatin administration (1 mg/kg, IP) from day 7 to day 10 and LPS injection (2 mg/kg, IP) on day 9. Blood was collected on day 10 for biochemical analysis. (**B**) Blood samples were analyzed for BUN and creatinine using a blood chemistry analyzer. \*\*\**P* < 0.001 vs. vehicle-treated control; *n* = 8–15. (**C**) Blood samples were analyzed for creatine kinase using a blood chemistry analyzer; *n* = 8. (**D**) Kidney sections were prepared from mice 48 h after LPS treatment and subjected to immunostaining for KIM-1. Representative images from six experiments show increased KIM-1 expression following combined atorvastatin (Atorva) and LPS treatment compared with the vehicle-treated control (CTL). Scale bars: 20 μm for the low-magnification images and 10 μm for the high-magnification image. (**E**) Kidney sections were prepared from mice 30 h after LPS treatment and subjected to immunostaining for NGAL. Representative images from four experiments show increased NGAL expression following atorvastatin and LPS treatment. Scale bars: 20 μm for the low-magnification images and 10 μm for the high-magnification image. Data in (**B** and **C**) are presented as mean ± SEM, with individual data points shown.

Because statins inhibit HMGCR and 25-HC suppresses SREBP-dependent feedback activation through INSIG proteins, we hypothesized that their combination would profoundly disrupt the mevalonate pathway leading to HK-2 cell toxicity (Fig. 1E, F) (22–24). In accordance with this, supplementation with either mevalonolactone or GGPP prevented HK-2 cell death induced by atorvastatin plus 25-HC (Fig. 1G, H). Since GGPP supports protein geranylgeranylation independently of cholesterol synthesis, its comparable rescue effect suggests that impaired prenylation, particularly geranylgeranylation, rather than reduced cholesterol synthesis, is the main driver of toxicity.

We next examined whether the concentrations of statins and 25-HC used in our experiments were sufficient to suppress the SREBP2-mediated restorative feedback loop in the mevalonate pathway in HK-2 cells, resulting in substantial defects in protein prenylation and cell toxicity. We therefore measured *HMGCR* mRNA, a well-established SREBP2 target gene (24), and SREBP2 cleavage, a key step required for SREBP2 activation, following co-treatment with statins and 25-HC. Statins increased HMGCR mRNA and SREBP2 cleavage as compensatory responses, whereas 25-HC abolished both effects (Fig. 1I–K). Together, these findings indicate that 25-HC blocks the SREBP2-mediated feedback response to statins, thereby intensifying mevalonate-pathway suppression, impairing protein prenylation, and promoting proximal tubular cell injury.

### Combined statin and low-dose LPS treatment induces AKI in mice

Based on these *in vitro* findings, we next examined the interaction of statins and 25-HC *in vivo* in the mouse kidney. 25-HC is synthesized and released from activated macrophages in infectious and inflammatory conditions (19, 25). LPS potently stimulates macrophages to produce 25-HC via toll-like receptor 4 (TLR4) activation (19, 26, 27). In our experiments, we chose a low dose of LPS (2 mg/kg) that activates macrophages without inducing AKI, thereby mimicking the inflammatory conditions under which statins may trigger AKI. We fed a cohort of mice a high-fat diet beginning 1 week prior to the first statin injection to enhance their inflammatory response to LPS (28) and to better mimic the metabolic conditions under which statins are commonly prescribed (Fig. 2A). After four daily intraperitoneal (IP) injections of atorvastatin (1 mg/kg) and a single IP injection of LPS (2 mg/kg), we measured blood urea nitrogen (BUN) and creatinine using a Dri-Chem 4000i dry chemistry analyzer (Fuji) (Fig. 2A). We found that the combination of atorvastatin and LPS, but neither atorvastatin nor LPS alone, increased plasma levels of BUN and creatinine, which are key biochemical indicators of AKI (Fig. 2B). Because rhabdomyolysis is a well-known adverse effect of statin therapy, and AKI can arise secondary to the massive release of myoglobin into the circulation during this process (29), we looked for possible elevations in creatine kinase (CK) that may indicate muscle injury. We did not find any indication that the combination of atorvastatin and LPS increased blood CK levels (Fig. 2C).

To determine whether acute tubular necrosis contributes to statin-induced AKI, we assessed the expression of the proximal tubular injury markers kidney injury molecule-1 (KIM-1) and neutrophil gelatinase-associated lipocalin (NGAL) in proximal tubular cells isolated from mice subjected to the same treatment regimen used for the BUN and creatinine measurements. We found that atorvastatin in combination with LPS induced the expression of both KIM-1 and NGAL in proximal tubular cells (Fig. 2D, E), consistent with the elevated BUN and creatinine levels observed in the same mice. We also examined the effects of fluvastatin (25 mg/kg/day) in combination with LPS (2 mg/kg) on NGAL and the apoptotic marker cleaved caspase-3. We found that the combination of fluvastatin and LPS induced widespread NGAL expression and, in a subset of kidney tubules, caused cleaved caspase-3-positive proximal tubular cells to be shed into the tubular lumen (Supplementary Fig. 3A, B). However, periodic acid-Schiff (PAS) staining revealed no discernible—or only minimal— histopathological abnormalities in the glomerular and tubular structures (Supplementary Fig. 3C), a finding known to occur in patients with AKI (17). These findings suggest that, under LPS-induced inflammatory conditions, statins can induce AKI by directly targeting renal tubular cells through interaction with 25-HC.

### Statin-induced AKI requires macrophages in LPS-treated mice

Cholesterol 25-hydroxylase (CH25H), the enzyme responsible for synthesizing 25-HC, is induced in macrophages under inflammatory conditions (25–27). Thus far, our results suggest that the *in vivo* effects of LPS in statin-treated mice may depend on macrophages if mediated by 25-HC. We first sought to examine whether the combined effects of statin and LPS could be recapitulated *in vitro* in the presence of macrophages. We found that LDH levels did not increase in HK-2 cells treated with fluvastatin and LPS, remaining comparable to those in untreated control cells (Fig. 3A). Then, we generated conditioned media (CM) from either LPS-stimulated or naïve RAW264.7 cells, a murine macrophage cell line, and transferred the CM to HK-2 cells (Fig. 3B, left). In contrast to direct LPS exposure, culture in CM derived from LPS-stimulated RAW264.7 cells rendered HK-2 cells susceptible to fluvastatin-induced toxicity (Fig. 3B, right). These results support a model in which LPS promotes statin-induced renal cell toxicity through macrophage-derived factors.

**Fig. 3.**
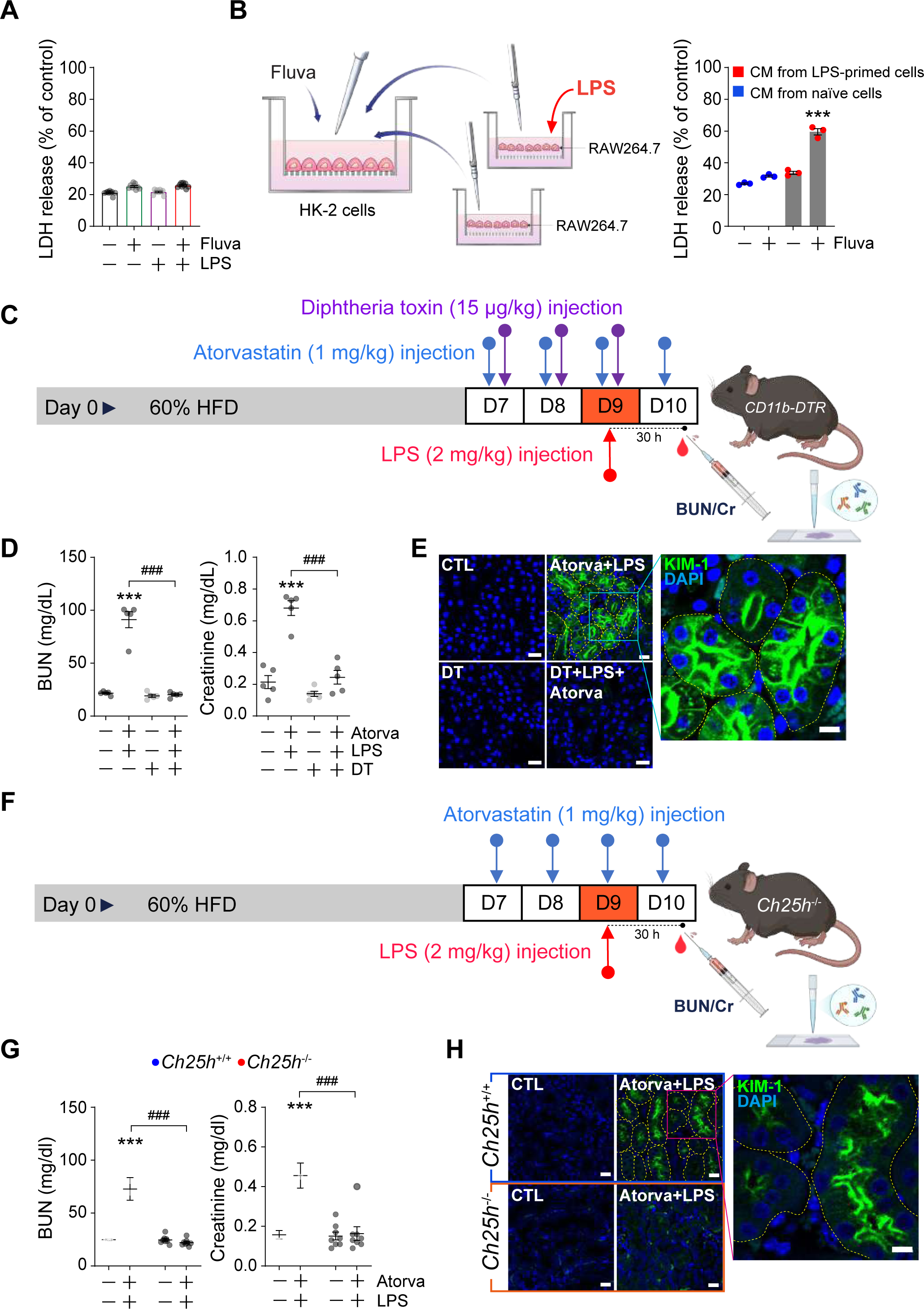
Activated macrophages are required for atorvastatin-induced AKI in LPS-treated mice. (**A**) Fluvastatin (Fluva, 0.5 μM) and/or LPS (2 μg/mL) were added to the HK-2 culture medium, and the cells were incubated for 72 h. Cell death was assessed using an LDH assay. (**B**) RAW264.7 cells were stimulated with LPS (2 μg/mL) or left untreated for 8 h, after which the conditioned media (CM) were collected and transferred to wells containing HK-2 cells. The HK-2 cells were then treated with fluvastatin (0.5 μM) for 72 h. \*\*\**P* < 0.001 vs. no-statin, no-priming control. *n* = 3. (**C**) Experimental scheme for AKI induction in B6.FVB-Tg(ITGAM-DTR/EGFP)34Lan/J mice with diphtheria toxin (DT, 15 μg/kg) administration. Mice fed a HFD received daily atorvastatin treatment (1 mg/kg) on days 7-10 and DT treatment during on days 7-9. LPS (2 mg/kg) was administered on day 9 in order to activate macrophages through TLR4 signaling. (**D**) Blood samples collected on day 10 were analyzed for BUN and creatinine levels using a blood chemistry analyzer. \*\*\**P* < 0.001 vs. vehicle-treated control; ^###^*P* < 0.001 vs. the atorvastatin plus LPS group. *n* = 5. (**E**) Kidney sections obtained 30 h after LPS treatment were subjected to immunofluorescence staining for KIM-1. Representative images from three experiments show DT prevents the induction of KIM-1 by atorvastatin and LPS. (**F**) Experimental scheme for AKI induction in *Ch25h^-/-^* mice. *Ch25h^+/+^* and *Ch25h^-/-^* mice fed a HFD received daily atorvastatin treatment (1 mg/kg) during days 7 to 10. LPS (2 mg/kg) was administered on day 9. (**G**) Blood samples collected on day 10 were analyzed for BUN and creatinine levels using a blood chemistry analyzer. \*\*\**P* < 0.001 vs. *Ch25h^+/+^* vehicle-treated control group; ^###^*P* < 0.001 vs. *Ch25h^-/-^* mice treated with atorvastatin and LPS. *n* = 8. (**H**) Kidney sections obtained 30 h after LPS treatment were subjected to immunofluorescence staining for KIM-1. Representative images from three experiments show atorvastatin combined with LPS fails to induce KIM-1 expression in *Ch25h^-/-^* mice. Scale bars: 20 μm for the low-magnification images and 10 μm for the high-magnification images in panels (E) and (H). Data in (**A, B, D,** and **G**) are presented as mean ± SEM, with individual data points shown.

Next, we used integrin alpha M–diphtheria toxin receptor (DTR) transgenic mice (B6.FVB-Tg(ITGAM-DTR/EGFP)34Lan/J) to transiently deplete macrophages/monocytes upon administration of diphtheria toxin (DT) (Fig. 3C) (30). Flow cytometry analysis confirmed that DT (15 μg/kg) markedly reduced peritoneal macrophages (Supplementary Fig. 4). We found that depletion of macrophages/monocytes abolished the increase in BUN and creatinine levels induced by co-administration of atorvastatin and LPS (Fig. 3D), as well as the upregulation of KIM-1 expression in renal proximal tubular cells (Fig. 3E). These findings indicate that activated macrophages are required for the interaction between atorvastatin and LPS in the induction of AKI *in vivo*. To further determine whether macrophage-derived 25-HC directly contributes to statin-induced AKI, we used mice deficient in *Ch25h*^⁻/⁻^ (Fig. 3F). In these knockout mice, co-administration of atorvastatin and LPS did not increase BUN or creatinine levels (Fig. 3G) and failed to induce KIM-1 expression in renal proximal tubular cells (Fig. 3H), confirming the essential roles of activated macrophages and macrophage-derived 25-HC in this process.

### Mevalonate rescues atorvastatin-induced AKI in LPS-injected mice

We next asked whether enhancing flux through the mevalonate pathway by supplementation with exogenous mevalonate could prevent AKI induced by the combination of atorvastatin and LPS. We administered mevalonolactone (10 mg/kg, IP) to mice once daily for four consecutive days, beginning 2 days before LPS (2 mg/kg) treatment (Fig. 4A). Thirty hours after LPS treatment, blood was collected to measure plasma BUN and creatinine levels.

**Fig. 4.**
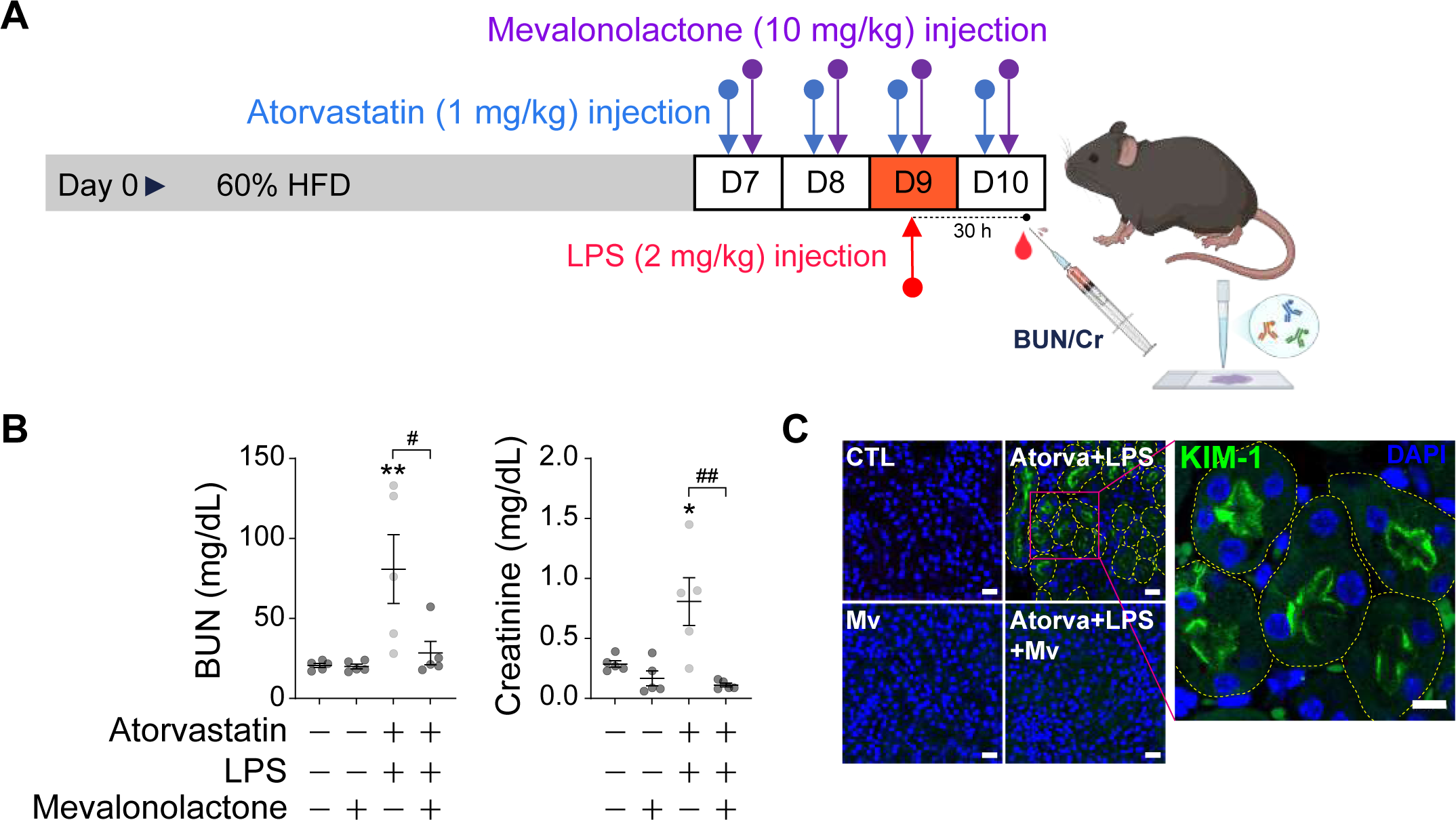
Mevalonolactone prevents atorvastatin-induced AKI in LPS-treated mice. (**A**) Schematic illustration of the AKI model and mevalonolactone treatment regimen. Mice fed an HFD were administered atorvastatin (1 mg/kg, IP) and mevalonolactone (10 mg/kg, IP) daily from day 7 to day 10. LPS (2 mg/kg, IP) was injected on day 9 to activate macrophages via TLR4 signaling. (**B**) Plasma BUN and creatinine levels were measured using a blood chemistry analyzer. Mevalonolactone significantly attenuated the increases in BUN and creatinine induced by combined atorvastatin and LPS treatment. \**P* < 0.05 and \*\**P* < 0.01 vs. untreated control; ^#^*P* < 0.05 and ^##^*P* < 0.01 for comparisons between groups with and without mevalonolactone in the presence of atorvastatin and LPS; *n* = 5. Data are presented as mean ± SEM, with individual data points shown. (**C**) Kidney sections collected 30 h after LPS treatment were subjected to immunofluorescence staining for KIM-1. Representative images from five experiments show mevalonolactone (Mv) prevents KIM-1 induction following combined atorvastatin and LPS treatment. Scale bars: 20 μm for the low-magnification images and 10 μm for the high-magnification image.

Mevalonolactone-treated mice did not show significant increases in BUN or creatinine in response to combined atorvastatin and LPS treatment (Fig. 4B). Consistently, mevalonolactone supplementation prevented the increase in KIM-1 expression in renal proximal tubules that was otherwise induced by the combination of atorvastatin and LPS (Fig. 4C). These data suggest that disruption of the mevalonate pathway contributes to atorvastatin-induced AKI in LPS-treated mice. Our *in vivo* AKI model thus recapitulates the injury-inducing interaction between statins and 25-HC observed *in vitro*, consistent with our findings in HK-2 cells treated with atorvastatin, 25-HC, and mevalonolactone.

### Combined atorvastatin and 25-HC treatment disrupts the autophagosome–lysosome pathway in proximal tubular cells

We next explored the molecular mechanism behind these observations. First, we examined possible changes in organelles such as mitochondria and lysosomes, which are particularly abundant in proximal tubular cells and play especially important roles in sustaining their high metabolic and homeostatic demands (31). Using transmission electron microscopy (TEM) imaging, we first identified alterations in the inner mitochondrial membrane ultrastructure, including numerous dilated and swollen cristae, in proximal tubule cells from mice treated with atorvastatin plus LPS (AKI group) compared with the other three experimental groups (Fig. 5A). We also observed the dramatic appearance of electron-dense lamellated structures (“lamellar bodies”, “zebra bodies”, or “myeloid bodies”) in the same AKI group (Fig. 5A; Supplementary Fig. 5A). Genetic conditions (such as Fabry disease and Sandhoff disease) or iatrogenic causes (such as drug-induced phospholipidosis) can lead to abnormal intracellular lamellar inclusions in various cell types, including renal cells, as a result of impaired lysosomal function (32–34). Thus, we asked whether autophagic clearance through the autophagosome–lysosome pathway might be impaired in the presence of atorvastatin and LPS. We first assessed the relevance of lamellar body formation to our AKI model using zoledronic acid, an inhibitor of FPP synthase (35). Zoledronic acid increased lamellar body formation in proximal tubular cells (Fig. 5B; Supplementary Fig. 5B), along with BUN and creatinine levels (Fig. 5C). These findings link impaired protein prenylation, which appears to be a key pathogenic mechanism in our AKI model, to lamellar body formation, providing a strong rationale for investigating autophagosome–lysosome dysfunction as a downstream mechanism.

**Fig. 5.**
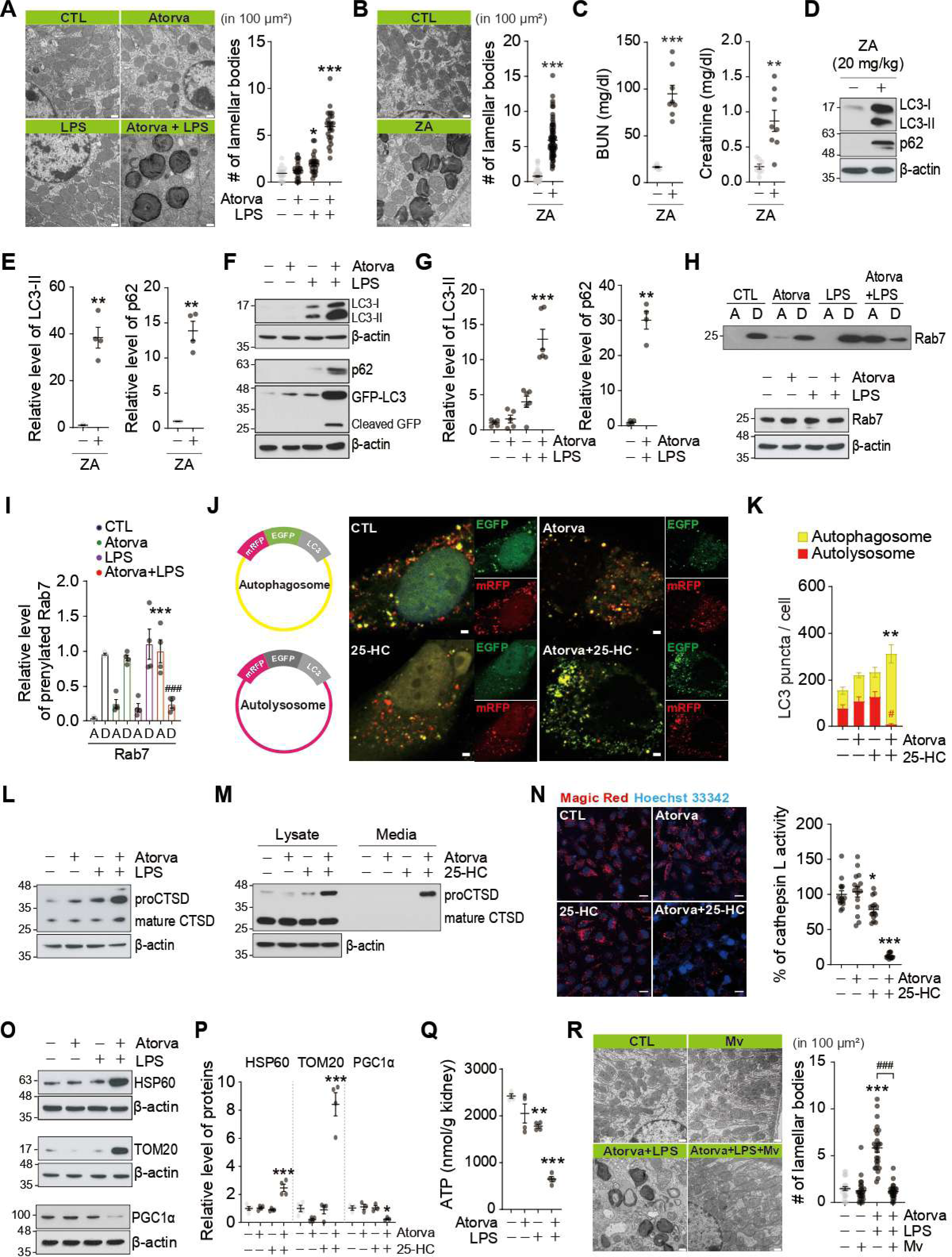
Atorvastatin combined with 25-HC induces dysfunction of the autophagosome– lysosomal pathway. (**A**) TEM images of proximal tubular cells from kidneys of mice treated with atorvastatin (1 mg/kg, IP) and/or LPS (2 mg/kg, IP). Kidneys were harvested 30 h after LPS administration and processed for TEM analysis. Combined treatment with atorvastatin and LPS induced the formation of lamellar bodies. The number of lamellar bodies per 100 μm² cell area was quantified and plotted. \**P* < 0.05 and \*\*\**P* < 0.001 vs. untreated control; *n* = 18-24 images from 3-4 mice per group. Scale bars: 500 nm. **(B)** TEM images of renal proximal tubular cells from mice treated with zoledronic acid (ZA; 20 mg/kg, intravenous) once daily for 2 consecutive days. Kidneys were collected 48 h after the final injection and processed for TEM analysis. \*\**P* < 0.001 vs. untreated control; n = 49–77 images from 4 mice per group. Scale bars, 500 nm. (C) Blood samples collected at the time of kidney harvest were analyzed for BUN and creatinine levels. \*\**P* < 0.01 and \*\*\**P* < 0.001 vs. untreated control; *n* = 8. (**D**) LC3 and p62 protein levels in kidney tissues were analyzed by Western blotting. ZA (20 mg/kg) treatment increased the levels of LC3-I, LC3-II, and p62 proteins. (**E**) Quantitative analysis of LC3-II and p62 protein band intensities shown in (D). \*\**P* < 0.01 vs. untreated control; *n* = 3-4. (**F**) Representative western blot analysis of kidney tissue showing increased accumulation of LC3-I, LC3-II, and p62 following combined atorvastatin (1 mg/kg) and LPS (2 mg/kg) treatment in GFP-LC3 mice. Cleaved GFP derived from GFP-LC3 was detected using an anti-GFP antibody. (**G**) Quantitative analysis of LC3-II and p62 protein band intensities shown in (F). \*\**P* < 0.01 and \*\*\**P* < 0.001 vs. untreated control; *n* = 4-6. (**H**) Rab7 prenylation was evaluated in kidney tissue using Triton X-114 phase partitioning. Representative blots show that combined atorvastatin and LPS treatment shifted Rab7 from the detergent-rich phase (D) to the aqueous phase (A) (upper panel), consistent with reduced Rab7 prenylation. Total Rab7 levels remained unchanged across treatment groups (lower panel). (**I**) Quantitative analysis of Rab7 prenylation. Values in the control group were normalized such that the aqueous and detergent-rich fractions summed to 1. \*\*\**P* < 0.001 vs. untreated control of aqueous phase; ^###^*P* < 0.001 vs. untreated control of detergent-rich phase; *n* = 4. (**J**) HK-2 cells were transfected with a tandem mRFP-EGFP-LC3 plasmid to evaluate autophagosome–lysosome fusion. Combined treatment with atorvastatin (5 μM) and 25-HC (0.1 μg/ml) increased the number of yellow puncta, representing autophagosomes, while decreasing the number of red puncta, representing autolysosomes. Scale bars: 2 μm. (**K**) Quantification of autophagosome and autolysosome puncta. \*\**P* < 0.01 vs. total puncta (yellow + red) in untreated control; ^#^*P* < 0.05 vs. red puncta in untreated control; *n* = 10. (**L**) Representative immunoblot images illustrating that combined treatment with atorvastatin (1 mg/kg) and LPS (2 mg/kg) increased proCTSD levels in mouse kidneys. Representative blot results from three independent experiments. (**M**) In HK-2 cells, combined treatment with atorvastatin (5 μM) and 25-HC (0.1 μg/ml) increased the levels of intracellular uncleaved proCTSD and secreted proCTSD. Representative blots from three independent experiments. (**N**) HK-2 cells treated with atorvastatin and 25-HC exhibited decreased fluorescence intensity of the Magic Red lysosomal cathepsin L activity dye. \**P* < 0.05 and \*\*\**P* < 0.001 vs. untreated control; *n* = 15 images from 3 independent experiments. Scale bars: 20 μm. (**O**) Typical western blot results showing that combined treatment with atorvastatin (1 mg/kg) and LPS (2 mg/kg) led to the accumulation of mitochondrial marker proteins, HSP60 and TOM20, in kidney tissue, while simultaneously reducing the levels of PGC1α, a master regulator of mitochondrial biogenesis. (**P**) Quantitative analysis of HSP60, TOM20, and PGC1α protein band intensities. \**P* < 0.05 and \*\*\**P* < 0.001 vs. untreated control; *n* = 3-4. (**Q**) ATP levels in kidney lysates were measured using an ATP assay kit. Co-treatment with atorvastatin and LPS significantly decreased ATP content in kidney tissue. \*\**P* < 0.01 and \*\*\**P* < 0.001 vs. untreated control; *n* = 4. (**R**) Mevalonolactone (Mv) suppressed lamellar body formation induced by combined atorvastatin and LPS treatment in mouse kidney proximal tubular cells. The number of lamellar bodies per 100 μm² of cell area was quantified and presented as a dot plot. \*\*\**P* < 0.001 vs. untreated control; ^###^*P* < 0.001 compared between mouse groups with and without mevalonolactone in the presence of atorvastatin and LPS; *n* = 24-25 images from 4 mice. Scale bars: 500 nm. Data in (**C, E, G, I, K, N, P, Q, and R**) are presented as mean ± SEM, with individual data points shown.

To determine whether blockade of prenylation disrupts the autophagosome–lysosome pathway, a critical cellular quality-control mechanism, we examined the effects of zoledronic acid on the accumulation of LC3-II and p62 (Fig. 5D). Zoledronic acid increased the levels of both LC3-II and p62 (Fig. 5D, E), indicating impaired autophagic flux and consequent accumulation of autophagosomes (36). Similarly, combined treatment with atorvastatin and LPS markedly increased both LC3-II and p62 levels in mice (Fig. 5F, G), further supporting impaired autophagic flux as a major defect induced by this combination. We next assessed LC3 processing in GFP-LC3 transgenic mice. In this model, GFP-LC3 cleavage occurs after delivery to the autolysosomal lumen, where the LC3 moiety is degraded more rapidly than GFP, resulting in the accumulation of free GFP (37). Notably, co-treatment with atorvastatin and LPS caused marked accumulation of both GFP-LC3 and cleaved free GFP (Fig. 5F), further supporting disruption of autophagic flux through the autophagosome–lysosome pathway.

Given the reduction in autophagic flux, we next investigated downstream processes, including autophagosome–lysosome fusion and lysosomal function. Because impaired prenylation appears to be the primary underlying defect in statin-induced AKI in this study, we initially focused on Rab7 as a candidate prenylated protein involved in the pathogenic mechanism, based on its established roles in both autophagosome–lysosome fusion and lysosomal function (38–40). First, we examined whether combined treatment with atorvastatin and LPS inhibits Rab7 prenylation using a Triton X-114 phase separation assay, in which non-prenylated proteins preferentially partition into the aqueous phase rather than the detergent phase (41). We found that the combined treatment increased the partitioning of Rab7 into the aqueous phase, suggesting inhibition of Rab7 prenylation (Fig. 5H, I). Next, we investigated whether autophagosome–lysosome fusion was impaired, given that Rab7 is a well-established tethering factor in this process (42). HK-2 cells were transfected with a tandem fluorescent LC3 construct (ptf-LC3; mRFP-EGFP-LC3), and the fluorescence shift from yellow puncta (autophagosomes) to red puncta (autolysosomes) was analyzed (43). Co-treatment with atorvastatin and 25-HC resulted in a marked increase in yellow puncta, in which both mRFP and EGFP signals were retained, along with a reduction in red puncta, where mRFP fluorescence remained dominant due to the quenching of EGFP in the acidic lysosomal environment. These findings indicate that autophagosome–lysosome fusion was impaired (Fig. 5J, K).

We also assessed lysosomal function by examining pro- and mature cathepsin D (CTSD) levels in mouse kidneys after co-administration of atorvastatin and LPS. Pro-CTSD is synthesized in the ER and, after transport to acidic late endosomes and lysosomes, is proteolytically cleaved into mature CTSD. Combined treatment increased pro-CTSD levels while leaving mature CTSD largely unchanged (Fig. 5L), thereby altering the immature-to-mature CTSD ratio. This pattern is commonly associated with moderate defects in lysosomal acidification or Rab7 knockdown (40, 44, 45). We also observed increased secretion of pro-CTSD into the culture medium of HK-2 cells treated with atorvastatin and 25-HC (Fig. 5M). Because Rab7 knockdown has been shown to increase secretion of pro-CTSD by impairing mannose-6-phosphate receptor recycling to the Golgi (40), the increased extracellular release of pro-CTSD in our setting may reflect defective cathepsin trafficking to lysosomes resulting from reduced Rab7 prenylation. To directly assess lysosomal enzyme function, we used the Magic Red fluorescent probe and found that combined treatment markedly reduced cathepsin L activity, indicating lysosomal dysfunction (Fig. 5N). Because cathepsin L contributes to CTSD processing (46), its reduced activity may partly explain the accumulation of pro-CTSD. Together, these findings suggest that atorvastatin and 25-HC impair not only autophagosome–lysosome fusion but also lysosomal function, potentially through defective Rab7 prenylation.

Mitochondria are energy-producing organelles that meet the high metabolic demands of the kidney. We investigated whether combined treatment impairs mitophagy. We examined changes in the levels of two key markers of mitochondrial mass, heat shock protein 60 (HSP60) and translocase of the outer mitochondrial membrane 20 (TOM20) (47). We found that both proteins accumulated in the kidneys of mice subjected to combined atorvastatin and LPS treatment, whereas peroxisome proliferator-activated receptor gamma coactivator 1-alpha (PGC-1α), a master regulator of mitochondrial biogenesis (48), was decreased (Fig. 5O, P). These findings indicate that mitophagy is impaired, leading to the accumulation of damaged mitochondria that would normally be removed to maintain mitochondrial quality control. Consistently, renal ATP levels were reduced after atorvastatin and LPS treatment (Fig. 5Q), supporting a role for mitochondrial dysfunction in this AKI model. Last, we tested whether supplementation of mevalonate, which rescues AKI phenotypes (see Fig. 4B, C), could inhibit the formation of lamellar bodies in mouse kidneys co-treated with atorvastatin and LPS, and found that it markedly reduced this effect (Fig. 5R).

### Atorvastatin induces AKI in a murine model of systemic infection

We next examined whether atorvastatin promotes AKI during infection, a condition that better recapitulates the clinical scenario than LPS exposure alone. To this end, mice fed a high-fat diet and treated with atorvastatin were infected with *Listeria monocytogenes* (10⁶ CFU) (Fig. 6A). We found that atorvastatin increased plasma BUN and creatinine levels in Listeria-infected mice, but not in non-infected controls (Fig. 6B). Immunofluorescence analysis further revealed increased NGAL expression in proximal tubular cells of infected mice receiving atorvastatin (Fig. 6C). Collectively, these findings suggest that infectious conditions enable atorvastatin to precipitate AKI.

**Fig. 6.**
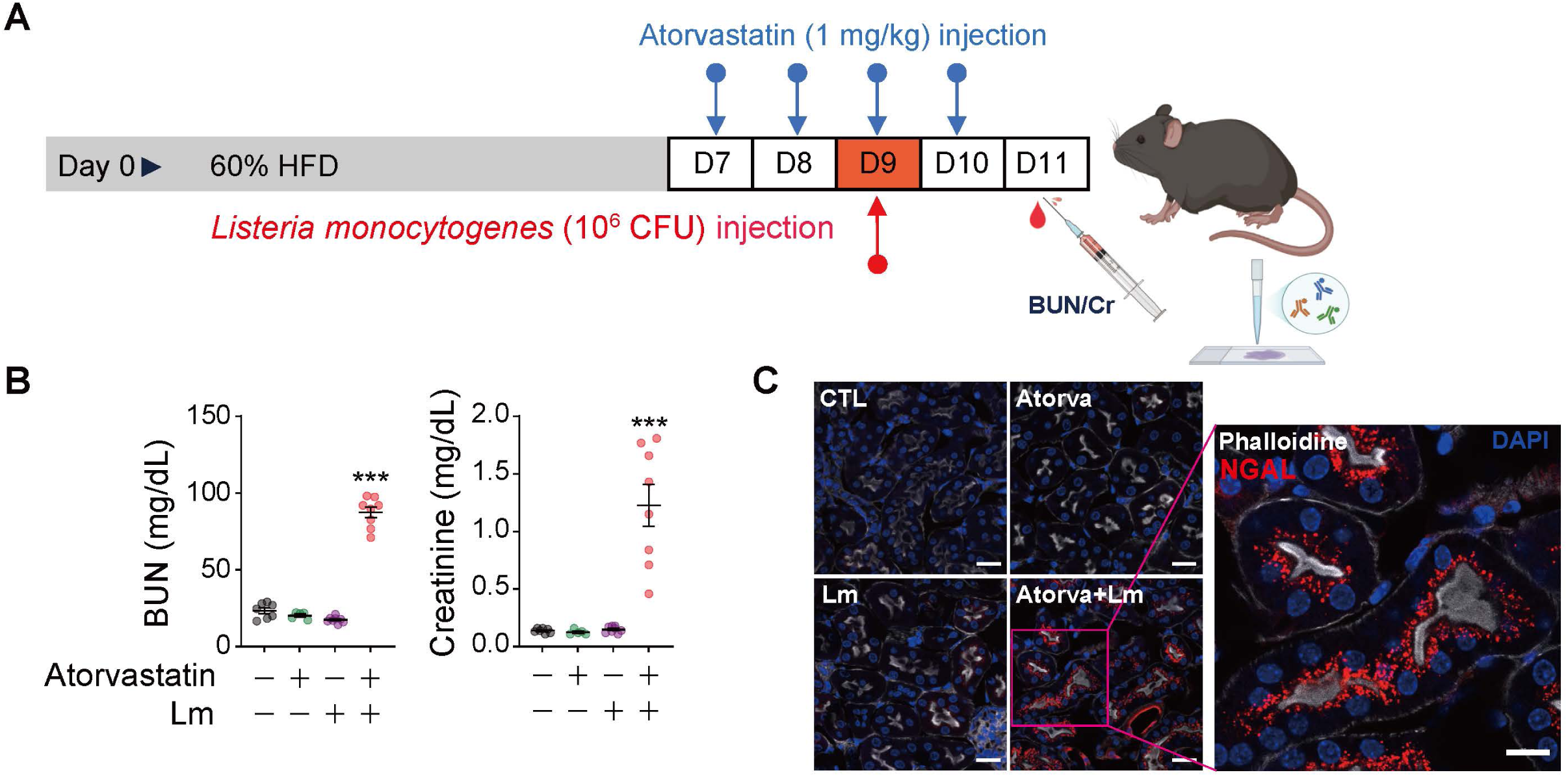
Atorvastatin induces AKI in mice infected with *L. monocytogenes*. (**A**) Mice maintained on a HFD received atorvastatin (1 mg/kg, IP) once daily from day 7 to day 10. On day 9, mice were challenged with 1 × 10⁶ CFU of *L. monocytogenes* (Lm). Blood and kidney samples were collected 48 h after bacterial infection. (**B**) Plasma BUN and creatinine levels at day 11 were measured using a blood chemistry analyzer. Atorvastatin significantly increased BUN and creatinine levels in Lm-infected mice. \*\*\**P* < 0.001 vs. untreated control. *n* = 5-8. Data are presented as mean ± SEM, with individual data points shown. (**C**) Kidney sections collected 48 h after bacterial infection were subjected to immunofluorescence staining for NGAL. Representative images from three independent experiments show that atorvastatin markedly increased NGAL expression in the kidneys of infected mice. Scale bars: 20 μm for the low-magnification images and 10 μm for the high-magnification image.

### Pexidartinib prevents AKI in mice co-treated with atorvastatin and LPS

We investigated the potential for pharmacological modulation of statin-induced AKI using a small molecule targeting key cell types involved in disease pathogenesis. Pexidartinib (PLX3397) is the first approved systemic therapy for patients with tenosynovial giant cell tumor (49). Its primary target is the colony-stimulating factor-1 receptor (CSF1R), which is expressed in myeloid lineage cells, including macrophages, monocytes, microglia, and osteoclasts (50). Pharmacological inhibition of CSF1R has been shown to effectively deplete macrophages (51, 52). Thus, pexidartinib (10 mg/kg) was administered intraperitoneally before statin and LPS treatment (Fig. 7A). We first confirmed the efficacy of pexidartinib in depleting macrophages by observing a reduction in the number of peritoneal macrophages (Supplementary Fig. 4). Notably, pexidartinib administration attenuated the elevations in BUN and creatinine observed in mice co-treated with atorvastatin and LPS (Fig. 7B). In addition, pexidartinib markedly decreased NGAL expression in statin/LPS co-treated mice (Fig. 7C). Together, these findings support a critical role for macrophages in atorvastatin-induced AKI and identify CSF1R targeting as a potential strategy to mitigate statin-associated renal toxicity.

**Fig. 7.**
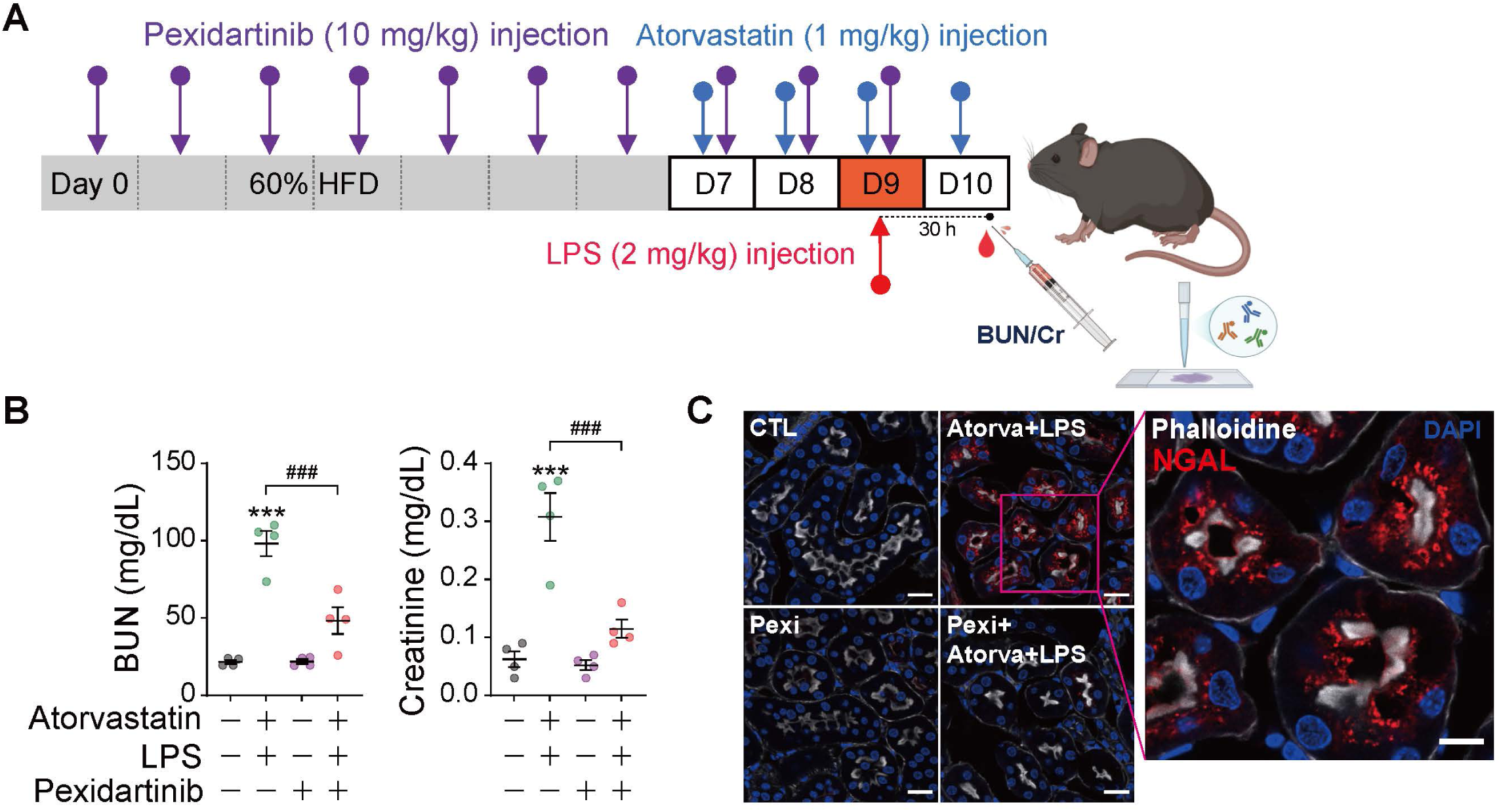
Pexidartinib attenuates atorvastatin-induced AKI in LPS-treated mice. (**A**) Mice were treated with pexidartinib (10 mg/kg, IP) daily from day 0 to day 9 while being maintained on a HFD. Atorvastatin (1 mg/kg, IP) was administered from day 7 to day 10. On day 9, mice received a single intraperitoneal injection of LPS (2 mg/kg). Blood and kidney tissues were collected 30 h after LPS injection. (**B**) Plasma BUN and creatinine levels were measured using a blood chemistry analyzer. Pexidartinib significantly attenuated the increases in BUN and creatinine levels induced by combined atorvastatin and LPS treatment. \*\*\**P* < 0.001 vs. untreated control; ^###^*P* < 0.001 for the comparison between atorvastatin- and LPS-treated mice with and without pexidartinib; *n* = 4. Data are presented as mean ± SEM, with individual data points shown. (**C**) Kidney sections collected 30 h after LPS treatment were subjected to immunofluorescence staining for NGAL. Representative images from three independent experiments show that pexidartinib (Pexi) inhibited NGAL induction caused by combined treatment with atorvastatin and LPS. Scale bars: 20 μm for the low-magnification images and 10 μm for the high-magnification image.

### Statin/25-HC co-treatment induces comparable pathological changes in human induced pluripotent stem cell (iPSC)-derived kidney organoids

Human iPSC-derived kidney organoids (53) were generated through 30 days of culture (Supplementary Fig. 6A) to determine whether this model could recapitulate the pathogenic interaction between statins and 25-HC. The resulting organoids comprised nephrin-positive glomerular cells, *Lotus tetragonolobus* lectin (LTL)-positive proximal tubular cells, and E-cadherin-positive distal tubular cells (Fig. 8A, Supplementary Fig. 6B). After treating the kidney organoids with fluvastatin (5 µM) in combination with 25-HC (0.1 μg/mL) for 72 h, we assessed proximal tubular cell injury by measuring KIM-1 expression. We found that neither fluvastatin nor 25-HC alone induced KIM-1 expression in any part of the kidney organoids, but the combination of fluvastatin and 25-HC induced KIM-1 expression almost exclusively in the LTL-positive proximal tubular cells of the organoids (Fig. 8A, B). These data demonstrate that fluvastatin induces proximal tubular cell injury in a 3D human kidney organoid model in the presence of 25-HC. We also examined the formation of lamellar bodies in kidney organoids using TEM. Kidney organoid cells co-treated with atorvastatin (5 µM) and 25-HC (0.1 μg/mL) showed an increased number of lamellar bodies in proximal tubular cells, which was reduced in the presence of mevalonolactone (Fig. 8C). Collectively, our results demonstrate that the renal pathological changes driven by the combined effects of statins and 25-HC are reproducible in human kidney organoids, highlighting the relevance of these pathological mechanisms to human kidney cells.

**Fig. 8.**
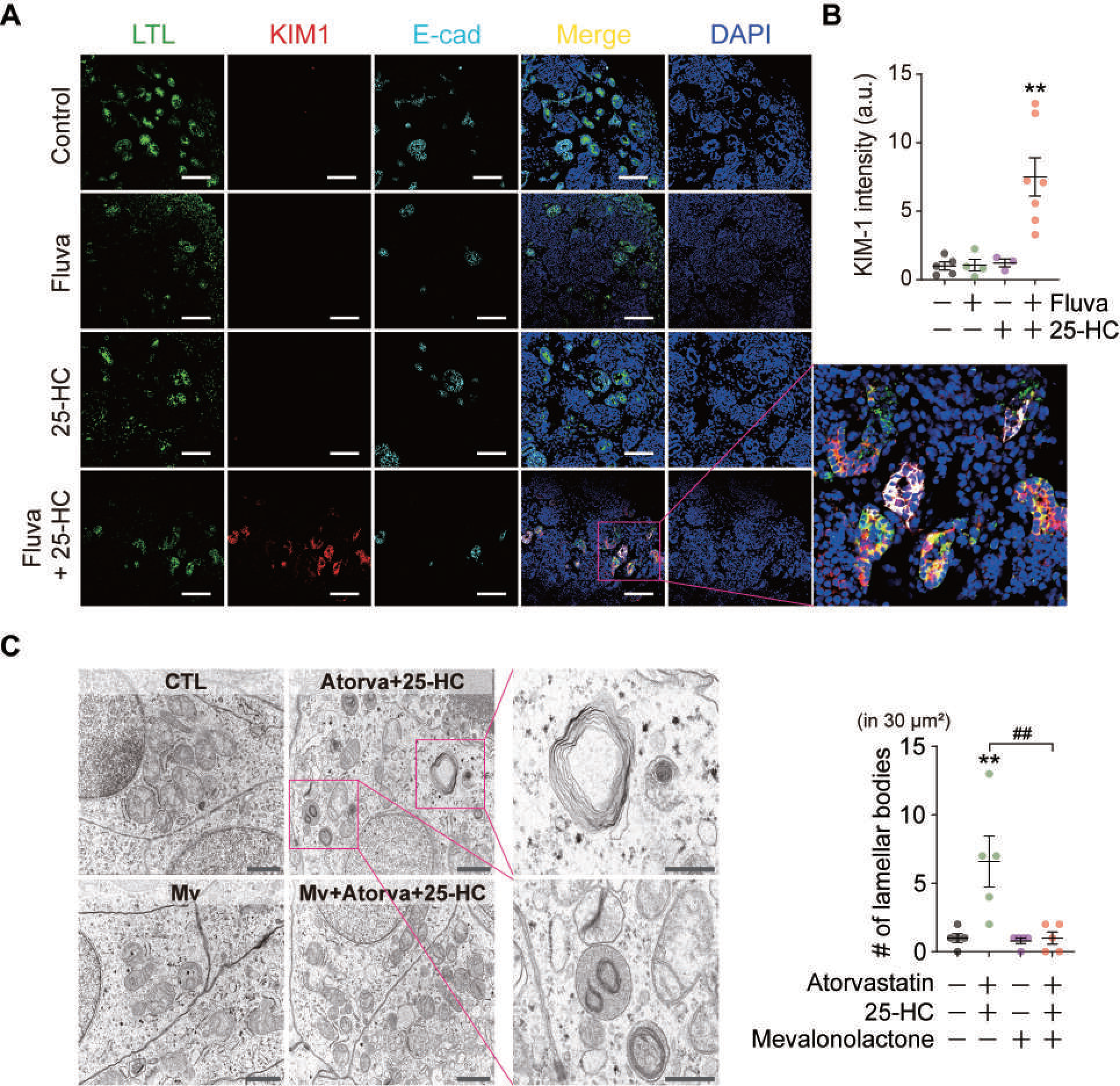
Combined statin and 25-HC treatment induces proximal tubular cell injury in kidney organoids. (**A**) Representative immunofluorescence images of kidney organoids treated with 25-HC (0.1 μg/mL) and/or fluvastatin (Fluva, 5 μM) for 72 h. Combined treatment markedly increased KIM-1 expression in proximal tubular cells compared with untreated controls or treatment with either fluvastatin or 25-HC alone. LTL and E-cadherin (E-cad) were used as markers of proximal and distal tubules, respectively. Scale bars: 100 μm. (**B**) Quantification of KIM-1 expression. Each dot represents the mean KIM-1 fluorescence intensity measured within proximal tubule regions of kidney organoids. \*\**P* < 0.01 vs. untreated control; *n* = 3–7. (**C**) Lamellar body formation visualized by TEM. The number of lamellar bodies per 30 μm² of cell area was quantified and presented as a dot plot. Mevalonolactone (Mv) inhibited lamellar body formation induced by the combined treatment with atorvastatin (Atorva, 5 µM) and 25-HC (0.1 μg/mL). \*\**P* < 0.01 vs. untreated control; ^##^*P* < 0.01 compared between groups with and without mevalonolactone in the presence of atorvastatin and 25-HC; *n* = 5. Scale bars: 1 μm for the low-magnification images and 500 nm for the high-magnification images. Data in (**B and C**) are presented as mean ± SEM, with individual data points shown.

## DISCUSSION

In our *in vitro* studies, we directly examined the combined effects of statins and 25-HC. By contrast, our *in vivo* experiments used LPS rather than exogenous 25-HC to model this interaction, because LPS stimulates macrophages to produce and release endogenous 25-HC. LPS, interferons, and viral infections are well known to induce the expression of CH25H, the enzyme that produces 25-HC in activated macrophages (25–27). Therefore, this strategy helps overcome a key limitation associated with exogenously administered 25-HC, namely the uncertainty regarding its effective *in vivo* concentration and the duration of its biological activity. Indeed, activated macrophages have been reported to persist and continuously produce 25-HC over extended periods (27). Furthermore, LPS administration has been shown to elevate plasma 25-HC levels in healthy human volunteers, suggesting similar mechanisms may also operate in humans (27). In our study, macrophages/monocytes, the principal cellular source of 25-HC, functioned as the key cellular mediators through which LPS promoted statin-induced renal tubular injury in both *in vitro* and *in vivo* models (see Fig. 3). Moreover, mevalonolactone exerted comparable protective effects against injury induced by statin plus LPS *in vivo* and by statin plus 25-HC *in vitro*, indicating that these two experimental conditions share a common mechanism (see Fig. 1 and Fig. 4). Together, these findings support the use of LPS, rather than exogenous 25-HC itself, as a relevant stimulus for modeling statin/25-HC-mediated tubular injury *in vivo*.

Induction of AKI in mice treated with zoledronic acid strongly supports the hypothesis that defects in protein prenylation underlie proximal tubular cell injury caused by the combined use of statins and LPS *in vivo*, as well as statins and 25-HC *in vitro*. Zoledronic acid, a potent bisphosphonate, inhibits FPP synthase and is widely used clinically for the treatment of osteoporosis, multiple myeloma, and bone metastases (54, 55). Its therapeutic effect is mediated through inhibition of prenylation of small GTPases in osteoclasts, leading to impaired differentiation and increased apoptosis of these cells (35). Importantly, AKI associated with bisphosphonate use is a well-recognized adverse effect, particularly with high-dose intravenous administration (56). Most documented cases are attributed to acute tubular necrosis, the most frequent cause of AKI (56). Accordingly, clinical guidelines recommend monitoring serum creatinine to ensure the safe use of zoledronic acid. These observations suggest that excessive inhibition of protein prenylation may represent one mechanism of nephrotoxicity in humans. An intriguing possibility is that statins and zoledronic acid share a common mechanism underlying these adverse effects. Whereas high-dose zoledronic acid alone can induce AKI, statins appear to require concomitant elevation of 25-HC to inhibit prenylation to a degree sufficient to elicit a similar effect.

Proximal tubules are particularly susceptible to various forms of cellular stress, including hypoxic and ischemic injury (57), owing to their high ATP requirements for intensive solute transport and their abundance of oxygen-consuming mitochondria. Accordingly, the preferential vulnerability of proximal tubular cells in our statin-associated AKI model may, at least in part, also reflect their intrinsically high metabolic and energetic demands. Mitochondrial dysfunction is also a well-recognized mechanism of drug-induced AKI, with agents such as aminoglycosides and antitumor drugs disrupting mitochondrial homeostasis through diverse mechanisms (58). Thus, mitochondrial dysfunction represents a plausible mechanism underlying statin-associated AKI. Given the distinct mechanistic features of our AKI model, in which the injury phenotype is largely attributable to defective protein prenylation, mitochondrial dysfunction, if it contributes substantially to the pathogenesis, would be expected to occur in close association with impaired prenylation. Consistent with this possibility, we proposed a model in which combined statin and 25-HC treatment impairs Rab7 prenylation, thereby disrupting autophagosome–lysosome function and mitophagy and ultimately compromising mitochondrial quality control. The result accumulation of dysfunctional mitochondria, together with cellular ATP depletion in renal proximal tubular cells, further supports the notion that the mitochondrial abnormalities observed in our study reflect functionally meaningful mitochondrial dysfunction arising in close association with defective protein prenylation. Together, these findings suggest that impaired prenylation may render proximal tubular cells particularly vulnerable by disrupting mitochondrial quality control and imposing an energetic deficit on cells with intrinsically high ATP demands, thereby providing a mechanistic basis for the preferential proximal tubular injury observed in statin-associated AKI.

The effectiveness of pexidartinib, a CSF1R inhibitor, together with the similar findings observed under DT-induced macrophage/monocyte depletion conditions in our murine statin-AKI model, strongly indicates that macrophages are a critical cellular component required for statin-induced AKI phenotypes. Moreover, the resistance to AKI induction observed in *Ch25h*^⁻/⁻^ mice supports our hypothesis that 25-HC produced in macrophages is a key mediator precipitating statin-induced kidney injury. Collectively, these findings raise the possibility that therapeutic strategies aimed at limiting macrophage-derived 25-HC production could mitigate susceptibility to statin-associated AKI. In this context, the protective effect of CSF1R inhibition provides proof of concept for targeting monocyte recruitment and differentiation as one potential therapeutic approach. This possibility is further supported by the high expression of CSF1R in circulating monocytes and monocyte-derived inflammatory macrophage subsets (50). In addition, therapeutic interventions designed to inhibit signaling pathways upstream of CH25H—for example, NF-κB inhibition by steroids or resveratrol— may be feasible (26, 59). Another potential strategy is to inhibit transmembrane receptors on monocytes and macrophages that play central roles in innate immune signaling, thereby preventing CH25H induction. TLR4 is one such key receptor, and its activation leads to robust and sustained upregulation of CH25H (26, 27). As TLR4 has emerged as a therapeutic target in multiple human diseases, numerous inhibitors have been evaluated in clinical trials (60). These efforts may ultimately create opportunities to develop agents capable of preventing statin-induced AKI.

## Ethics declaration

All animal protocols were approved (No. 2023-0150) by the Institutional Animal Care and Use Committee at Yonsei University College of Medicine under NIH guidelines.

## Author contributions

CHK and BSK conceptualized the study. HJW, EJK, YC, JHL, JY, HK, JM, YJ, HL and HYC performed the investigation. HJW, EJK, JHL, JY, HK and CHK prepared the figures. CHK, BSK, JL and HYG supervised the study. CHK and BSK wrote the original draft of the manuscript. CHK, BSK and HYG reviewed and edited the manuscript.

